# DeepPVP: phenotype-based prioritization of causative variants using deep learning

**DOI:** 10.1101/311621

**Authors:** Imane Boudellioua, Maxat Kulmanov, Paul N Schofield, Georgios V Gkoutos, Robert Hoehndorf

**Affiliations:** Computational Bioscience Research Center (CBRC), King Abdullah University of Science and Technology, 4700 KAUST, 23955-6900 Thuwal, Kingdom of Saudi Arabia.; Computer, Electrical and Mathematical Sciences & Engineering Division (CEMSE), King Abdullah University of Science and Technology, 4700 KAUST, PO Box 2882, 23955-6900 Thuwal, Kingdom of Saudi Arabia.; Department of Physiology, Development & Neuroscience, University of Cambridge, Downing Street, CB2 3EG Cambridge, United Kingdom.; College of Medical and Dental Sciences, Institute of Cancer and Genomic Sciences, Centre for Computational Biology, University of Birmingham, B15 2TT Birmingham, United Kingdom.; Institute of Translational Medicine, University Hospitals Birmingham, NHS Foundation Trust, B15 2TT Birmingham, United Kingdom.; NIHR Experimental Cancer Medicine Centre, B15 2TT Birmingham, United Kingdom.; NIHR Surgical Reconstruction and Microbiology, B15 2TT Birmingham, United Kingdom.; NIHR Biomedical Research Centre, B15 2TT Birmingham, United Kingdom.; MRC Health Data Research UK, B15 2TT Birmingham, United Kingdom.

**Keywords:** variant prioritization, phenotype, machine learning, ontology

## Abstract

**Background:** Prioritization of variants in personal genomic data is a major challenge. Recently, computational methods that rely on comparing phenotype similarity have shown to be useful to identify causative variants. In these methods, pathogenicity prediction is combined with a semantic similarity measure to prioritize not only variants that are likely to be dysfunctional but those that are likely involved in the pathogenesis of a patient’s phenotype.

**Results:** We have developed DeepPVP, a variant prioritization method that combined automated inference with deep neural networks to identify the likely causative variants in whole exome or whole genome sequence data. We demonstrate that DeepPVP performs significantly better than existing methods, including phenotype-based methods that use similar features. DeepPVP is freely available at https://github.com/bio-ontology-research-group/phenomenet-vp.

**Conclusions:** DeepPVP further improves on existing variant prioritization methods both in terms of speed as well as accuracy.

## Background

There is now a large number of methods available for prioritizing variants in whole exome or whole genome datasets [1]. These methods commonly identify the variants which are pathogenic, i.e., the variants that may alter normal functions of a protein, either directly through a change in a protein’s amino acid sequence or indirectly through a change of expression [2, 3, 4]. In coding and noncoding DNA sequences, there are usually multiple variants that could possibly be pathogenic, but most of them are sub-clinical or will not result in any detectable phenotypic manifestations [5].

Recently, several methods have become available that utilize information about phenotypes observed in a patient to identify potentially causative variants [6, 7, 8, 9]. Phenotypes are useful for identifying gene–disease associations because they implicitly reflect interactions occurring within an organism across multiple levels of organisation [10, 11, 12]. Phenotype-based methods work by comparing the phenotypes of a patient with a knowledgebase of gene-to-phenotype associations. A measure of phenotypic similarity is computed between a patient’s phenotypes and abnormal phenotypes associated with gene variants or mutations. The phenotypic similarity is then used either as a filter to remove pathogenic variants in genes that are not associated with similar phenotypes to the ones observed in the patient [9] or as a feature in machine learning approaches [6, 7].

The gene-to-phenotype associations used in phenotype-based prioritization strategies come from clinical observations such as those reported in the Online Mendelian Inheritance in Man (OMIM) database [13] or in the ClinVar database [14]. In some cases, they may also come from model organisms. Comparing model organism phenotypes to human phenotypes (i.e., the phenotypes observed in a patient) requires a framework that allows phenotypes of different species to be compared, such as the Uberpheno [15] or PhenomeNET ontology [16].

We have previously developed the PhenomeNET Variant Predictor (PVP) [7] to prioritize causative variants in personal genomic data. We have shown that PVP outperforms other phenotype-based approaches such as the Exomiser or Genomiser tools [17, 18], or Phevor [9]. PVP is based on a random forest classifier, similarly to Exomiser and Genomiser, which also use a random forest. Features used to classify a variant as causative or non-causative combine a phenotype similarity score (to prioritize a gene as being associated with the phenotypes observed in the patient) and a pathogenicity score, as well as other features such as the mode of inheritance and genotype of the variant. As most variants are neutral, there is a very large imbalance between positive and negatives, and the challenges for building a machine learning model for finding causative variants is to account for this imbalance during training and testing.

Recently, deep neural networks have shown to be successful in many domains [19] and often result in better classification performance [4]. We have developed DeepPVP, an extension of the PVP system which uses deep learning and achieves significantly better performance in predicting disease-associated variants than the previous PVP system, as well as competing algorithms that combine pathogenicity and phenotype similarity. DeepPVP not only uses a deep artificial neural network to classify variants into causative and non-causative but also corrects for a common bias in variant prioritization methods [20, 21] in which gene-based features are repeated and potentially lead to overfitting. DeepPVP is freely available at https://github.com/bio-ontology-research-group/phenomenet-vp.

## Implementation

### Training and testing data

We downloaded the ClinVar database 7th Feb, 2017, and extracted GRCh37 genomic variants annotated with at least one disease from OMIM, characterized as *Pathogenic* in their clinical significance, and not annotated with *conflicting interpretation* in their review status. We obtained 31,156 pathogenic variants associates with 3,938 diseases in total and the set of these variants constitutes candidate positive instances in our training dataset. We also extracted GRCh37 genomic variants that are characterized as *Benign* in clinical significance, and not annotated with *conflicting interpretation* in their review status. We obtained 23,808 such benign genomic variants from ClinVar which form the candidate negative instances in our training dataset. We excluded any variant records mapped to more than one gene and variant records with missing information on the reference or alternate alleles. For pathogenic variant records, we defined variant-disease pairs for each pathogenic variant and its associated OMIM disease. In our dataset, some pathogenic variants are annotated with multiple OMIM diseases. For each of these variants and the *n* OMIM diseases they may cause, we created *n* variant-disease pairs. For example, variant *rs201108965* in *TMEM216* is annotated with two diseases; Joubert syndrome 2 (`OMIM:608091`) and Meckel syndrome type 2 (`OMIM:603194`). We define two variant-disease pairs (*rs201108965*, `OMIM:608091`) and (*rs201108965*, `OMIM:603194`). After this step, we have 30,770 pathogenic variant-disease pairs and 20,174 benign variants.

In DeepPVP, we use the zygosity of a variant as one of the training features. The zygosity information is not provided in ClinVar. In a given Variant Call Format (VCF) [22] file, zygosity is represented in the genotype field. For instance, a heterozygous variant will have a genotype value `0/1`, while a homozygous variant will have a genotype value `1/1` in the VCF file. We assigned the genotype information to our pathogenic variant-disease pairs based on the mode of inheritance associated with the disease caused by the variant. We extracted the mode of inheritance of the associated OMIM disease using the information provided by the HPO annotations of OMIM diseases [23]. If the disease’s mode of inheritance is recessive, we assigned the zygosity of the variant as homozygote (denoted with genotype `1/1`). In this case, we created a variant-disease-zygosity triple representing this information. If the OMIM disease’s mode of inheritance is not recessive (i.e., any other mode of inheritance, including dominant, unknown, X-linked, etc.), we generated two variant-disease-zygosity triples and and characterize one of them as homozygote (denoted with genotype `1/1`) and another as heterozygote (denoted with genotype `0/1`). For example, pathogenic variant *rs397704705* in *AP5Z1* is associated with Spastic paraplegia 48 (`OMIM:613647`). This OMIM disease is recessive and, hence, we characterize variant *rs397704705* with genotype `1/1`, generating a variant-disease-zygosity triple consisting of variant *rs397704705*, disease `OMIM:613647`, and genotype `1/1`. Another example is the pathogenic variant *rs387907031* in *ARHGAP31* associated with Adams-Oliver syndrome 1 (`OMIM:100300`). This disease is dominant and, hence, we generated two variant-disease-zygosity triples: variant *rs387907031*, disease `OMIM:100300`, and the genotype `0/1`, and variant *rs387907031*, disease `OMIM:100300`, and genotype `1/1`. Since benign variants are not associated with a disease or mode of inheritance, we treat each of them as both a homozygote and heterozygote, generating two variant-zygosity pairs for each benign variant. After this step, we obtained 61,540 triples consisting of pathogenic variant, disease, and zygosity, and 40,348 pairs of benign variant and zygosity.

The triples consisting of variant, disease, and zygosity constitute positive samples. For each positive instance (*V, D, Z*) consisting of a variant, disease, and zygosity, we randomly select, with equal probability, one of two possible negative instances: a randomly selected benign variant in the same gene as *V*, or a triple (*V, D′, Z*) where *D*′ ≠ *D*. To map intergenic variants to genes, we link variants to their nearest gene.

For example, a positive instance in our training data is a pathogenic variant *rs267606829* in *FOXRED1*, associated with Mitochondrial complex I deficiency (`OMIM:252010`), as a homozygote. A negative instance according to the first strategy could be a benign variant, such as *rs1786702*, in *FOXRED1*, as a heterozygote. A negative instance according to the second strategy is the same pathogenic variant, *rs267606829* in FOXRED1, as a homozygote, but associated with another OMIM disease such as Tooth Agenesis (`OMIM:604625`).

As an independent and unseen evaluation dataset, we downloaded all variants from ClinVar released between Feb 8th 2017 and Jan 27th 2018. In this dataset, we processed all GRCh37 variants in the same manner as for our training dataset to construct triples of pathogenic variants, disease, and zygosity. However, if the OMIM disease’s mode of inheritance of the variant is not recessive (i.e., any other mode of inheritance, including dominant, unknown, X-linked, etc.), we assigned the zygosity randomly either as homozygote (denoted with genotype `1/1`), or heterozygote (denoted with genotype `0/1`). We obtained a total of 5,686 such triples associated with 1,370 diseases for validation.

### Generation of synthetic patients

In our evaluation, we generated a set of synthetic patients as a realistic evaluation case, similarly to previous work [18, 7]. We randomly selected a whole exome from the 1,000 Genomes project [24] and inserted a pathogenic variant *V*, assign the disease associated with *V* in ClinVar to the exome, and present *V* as a homo- or heterozygote based on the mode of inheritance associated with the disease. Each of these exomes together with the disease’s phenotypes and mode of inheritance form a synthetic patient in which we aim to recover the inserted variant.

### Annotating variants

Annotating variants with pathogenicy scores from CADD, DANN, and GWAVA is a time-consuming process in PVP [7] and other phenotype-based variant prioritization tools [18], especially when analyzing WGS data comprised of millions of variants. PVP 1.0 uses tabix [25] for indexing and retrieval of the pathogenicty scores per chromosome and genomic position. To optimize the annotation phase of DeepPVP, we extracted 31,491,995 variants from samples of the 1000 Genomes Project [24] and annotated them with pathogenicity scores from CADD, DANN, and GWAVA. DeepPVP keeps this set of pre-annotated variants in memory to provide fast retrieval of annotations for common variants. DeepPVP utilizes tabix only when the variant annotated is not available in the pre-annotated library, and therefore minimizes disk access.

### Model and availability

We implemented our DeepPVP deep neural network in Python 2.7. We used Keras [26] with a TensorFlow backend [27]. We used one hot encoding to represent our categorical feature of the inheritance mode of the disease. We handled missing values for CADD, GWAVA, DANN, and semantic similarity scores by mean imputation. We also added additional flags for missing values as features. We retrieved gene-phenotype association data from human and model organisms (mouse and zebrafish) on Feb 7th, 2017 and used them to generate the ontology and high level phenotypes and semantic similarity score features. We used the scikit-learn [28] library for tuning the hyperparameters of the neural network using grid search. We designed a sequential model with an input layer, three hidden layers of 67, 134, 67 neurons respectively with Rectified Linear Units (ReLU) [29] activation function, and an output layer with a sigmoid activation function. We trained the model using the Adam optimization algorithm [30] which has been widely adopted for deep learning as a computationally efficient, fast convergent, extension to stochastic gradient descent. We used dropout [31] between the hidden layers and the output layer to prevent overfitting. We trained our DeepPVP model for 300 epochs, 20,000 batch size, and a learning rate of 0.001. In training, we specified a 20% random validation, and monitored the validation loss with each epoch. We kept the rest of the parameters in their default values.

The DeepPVP system, the synthetic genome sequences and our analysis results can be found at https://github.com/bio-ontology-research-group/phenomenet-vp.

## Results

### DeepPVP: phenotype-based prediction using a deep artificial neural networks

We developed the Deep PhenomeNET Variant Predictor (DeepPVP) as a system to identify causative variants for patients based on personal genomic data as well as phenotypes observed in the patient. We consider a variant to be causative for a disease *D* if the variant is both pathogenic and affects a structure or function that leads to *D*. This distinction is motivated by the observation that healthy individuals can have multiple highly pathogenic variants resulting in a complete loss of function; it is therefore not usually sufficient to identify pathogenic variants alone as there may be many.

DeepPVP is a command-line tool which takes a Variant Call Format (VCF) [22] file as an input together with a set of phenotypes coded either through the Human Phenotype Ontology (HPO) [32] or the Mammalian Phenotype Ontology (MP) [33]. It outputs a prediction score for each variant in the VCF file; the prediction score measures the likelihood that a variant is causative for the phenotypes specified as input to the method.

To predict whether a variant is causative or not, DeepPVP uses similar features as the PVP system [7] and combines multiple pathogenicity prediction scores, a phenotype similarity computed by the PhenomeNET system, and a high-level phenotypic characterization of a patient. The full list of features used by DeepPVP is listed in Supplementary Table 1. All features can be generated from a patient’s VCF file and a set of phenotypes coded either with HPO or MP.

In DeepPVP, we use a deep neural network to classify variants as causative or non-causative. Specifically, DeepPVP uses a feed forward neural network with five layers (see Figure 1). The input layer in our architecture consists of 67 neurons (for the 67 features) and an output layer consisting of a single output neuron which outputs the prediction score of DeepPVP. DeepPVP uses three hidden layers with 67, 134, and 67 neurons, respectively. Each hidden layer uses a Rectified Linear Unit (ReLU) [29] activation function, and the output layer uses a sigmoid activation function.

**Figure 1:**
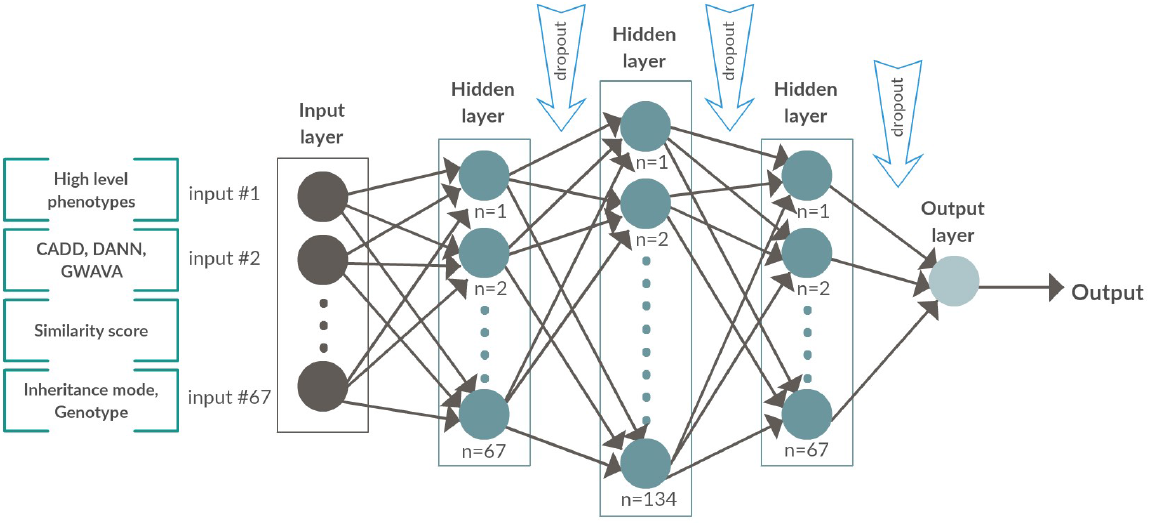
Overview over the DeepPVP neural network model.

DeepPVP is trained similarly as PVP to improve performance of identifying causative variants in real genomic sequences (in contrast to performance on a testing set). When training DeepPVP, we use as positive instances all causative variants from our training set together with the phenotypes of the disease for which they are causative. We discriminate these from two kinds of negatives: benign variants (i.e., variants that do not alter protein function) and pathogenic but non-causative variants. We consider pathogenic non-causative variants as pathogenic variants (in our training set) which are not associated with phenotypes of the disease they cause, but rather with a different disease. The aim of this selection strategy is to discriminate causative variants from all other variants.

We train the DeepPVP model using back-propagation, evaluate the model’s results on predicting causative variants, and compare against several competing methods. While the different evaluation scenarios omit some parts of the information about variants and the diseases they are associated with in order to not bias the evaluation results, we finally retrain a model using all available information and make it available as the final DeepPVP prediction model.

### Evaluating DeepPVP’s ability to find causative variants

We evaluate the performance of DeepPVP during training using both the accuracy and the binary cross entropy loss observed on the training and the validation set. The validation set consists of a randomly selected 20% set of variants from the training set. Figure 2 illustrates the results of this evaluation. We observe that both the training loss and validation loss decrease with the number of epochs; both training and validation loss as well as accuracy stabilize after over 100 epochs and are stable before we end the training after 300 epochs.

**Figure 2:**
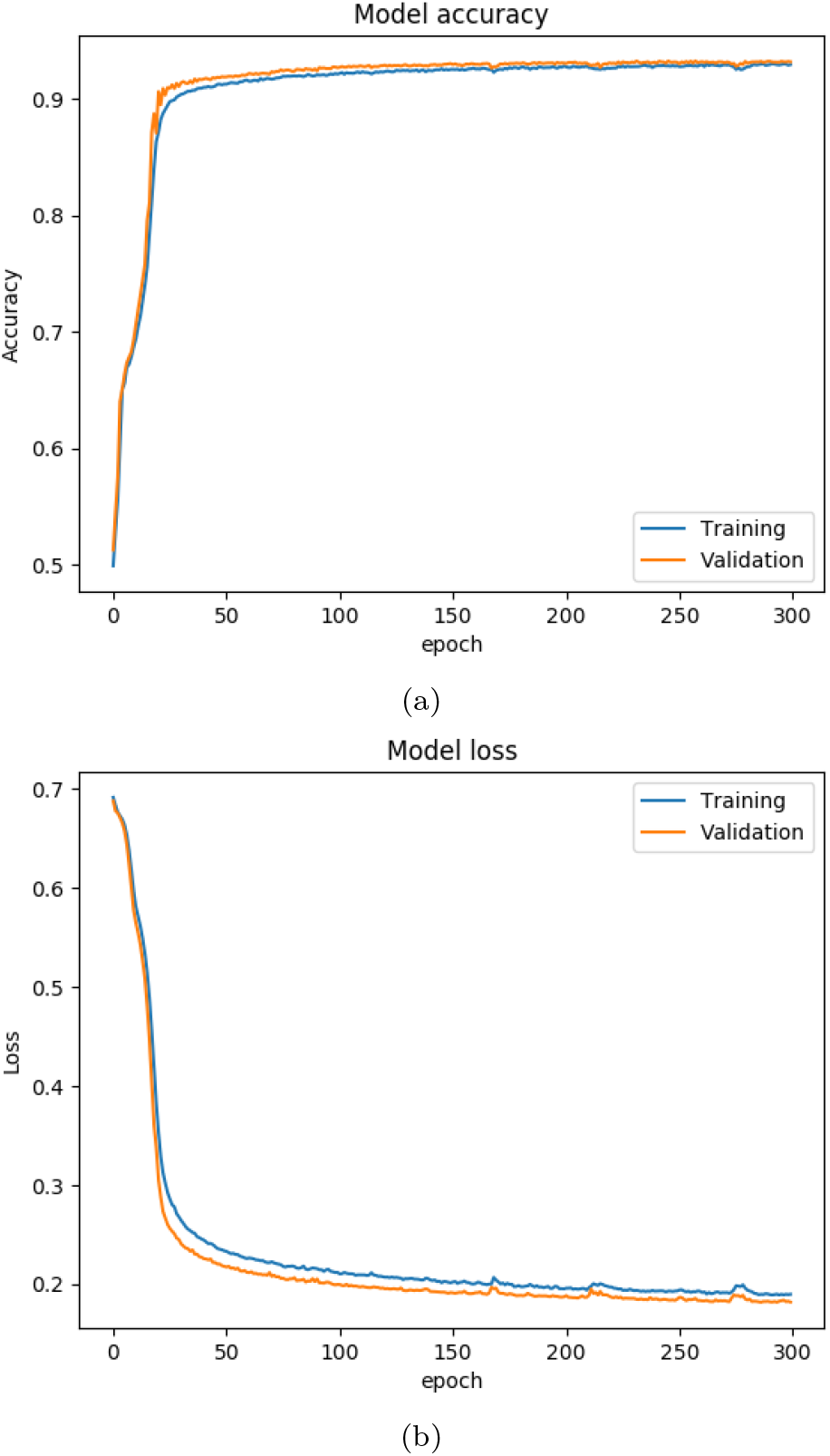
Performance of DeepPVP on convergence on training data and validati data (20% split)

We evaluate the ability of DeepPVP to predict causative variants using several approaches. First, we test the predictive performance of DeepPVP on a validation set consisting of a randomly selected 20% set of variants from the training set. Under this condition, the training and validation set will contain distinct variants, but the variants in training and test may be associated with the same disease. Figure 3 illustrates the receiver operating characteristic (ROC) curve [34] and precision-recall curve for this evaluation. We find that DeepPVP achieves an overall ROCAUC of 0.979, and AUPR of 0.977.

**Figure 3:**
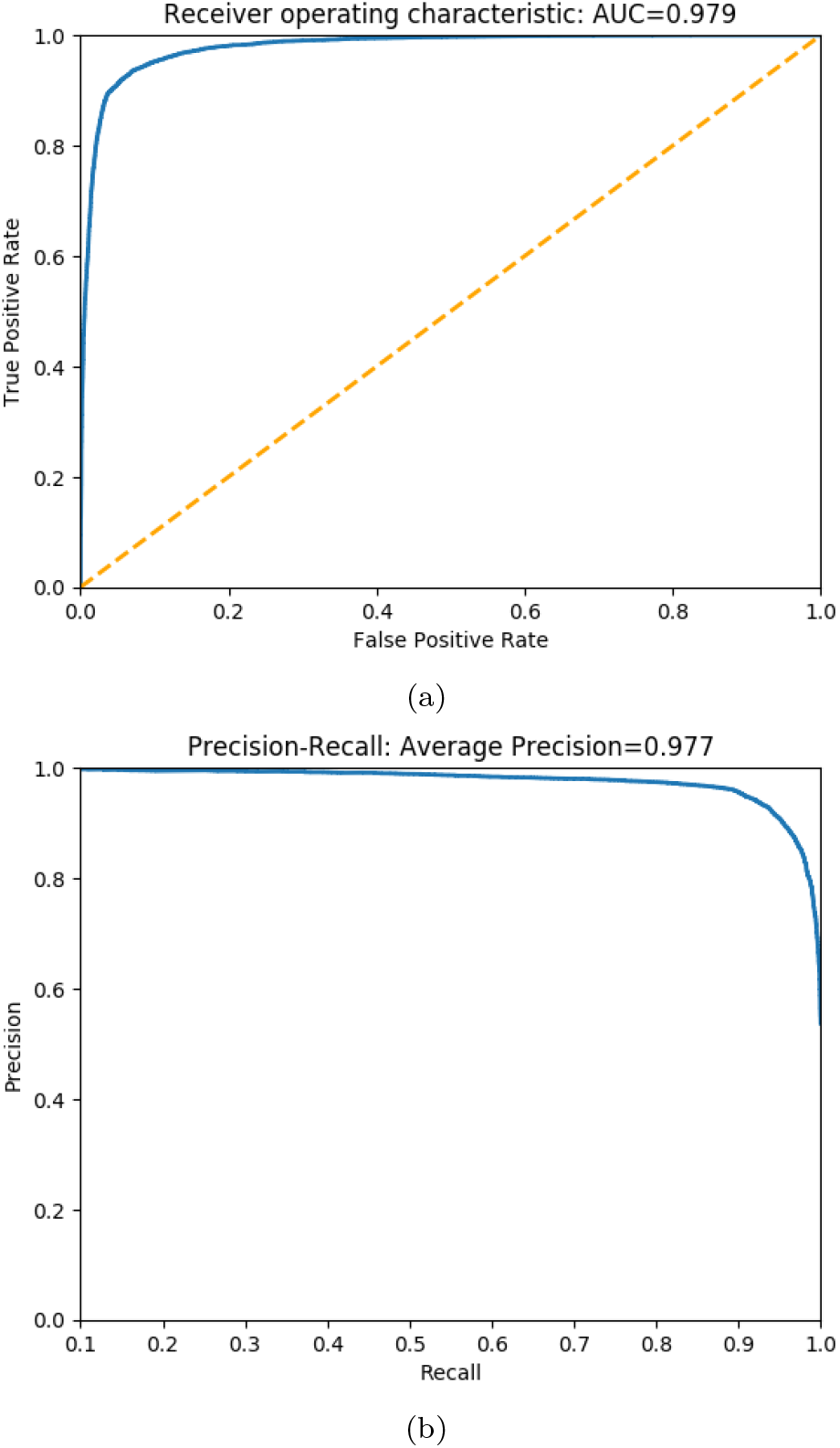
Performance of DeepPVP on a random test set (20% split)

Second, we test how well DeepPVP is able to find variants for diseases which have not been used in the training of DeepPVP at all. We use this strategy to determine DeepPVP’s performance on orphan diseases without any prior knowledge of associated genes. For this evaluation, we split our variant–disease pairs into a training and testing set. We stratified the two sets by disease so that we train on 75% of instances in our ClinVar-derived dataset (comprising 62% of diseases) and test on the remaining 25% of instances (and their diseases). Stratification by disease, in contrast to stratification by disease-variant pair, ensures that no disease-associated information (i.e., the phenotype scores between a disease and a gene, which would be identical for multiple disease-causing variants in the same gene) was seen during training and therefore unfairly bias the evaluation results. Figure 4 shows the ROC curve and the precision-recall curve in this evaluation. The area under the ROC curve (ROCAUC) is 0.943 and the area under the precision-recall curve is 0.921. The drop in predictive performance when stratifying training and testing data by disease compared to random split illustrates the performance that is to be expected for variants in diseases without prior knowledge about disease-associated genes.

**Figure 4:**
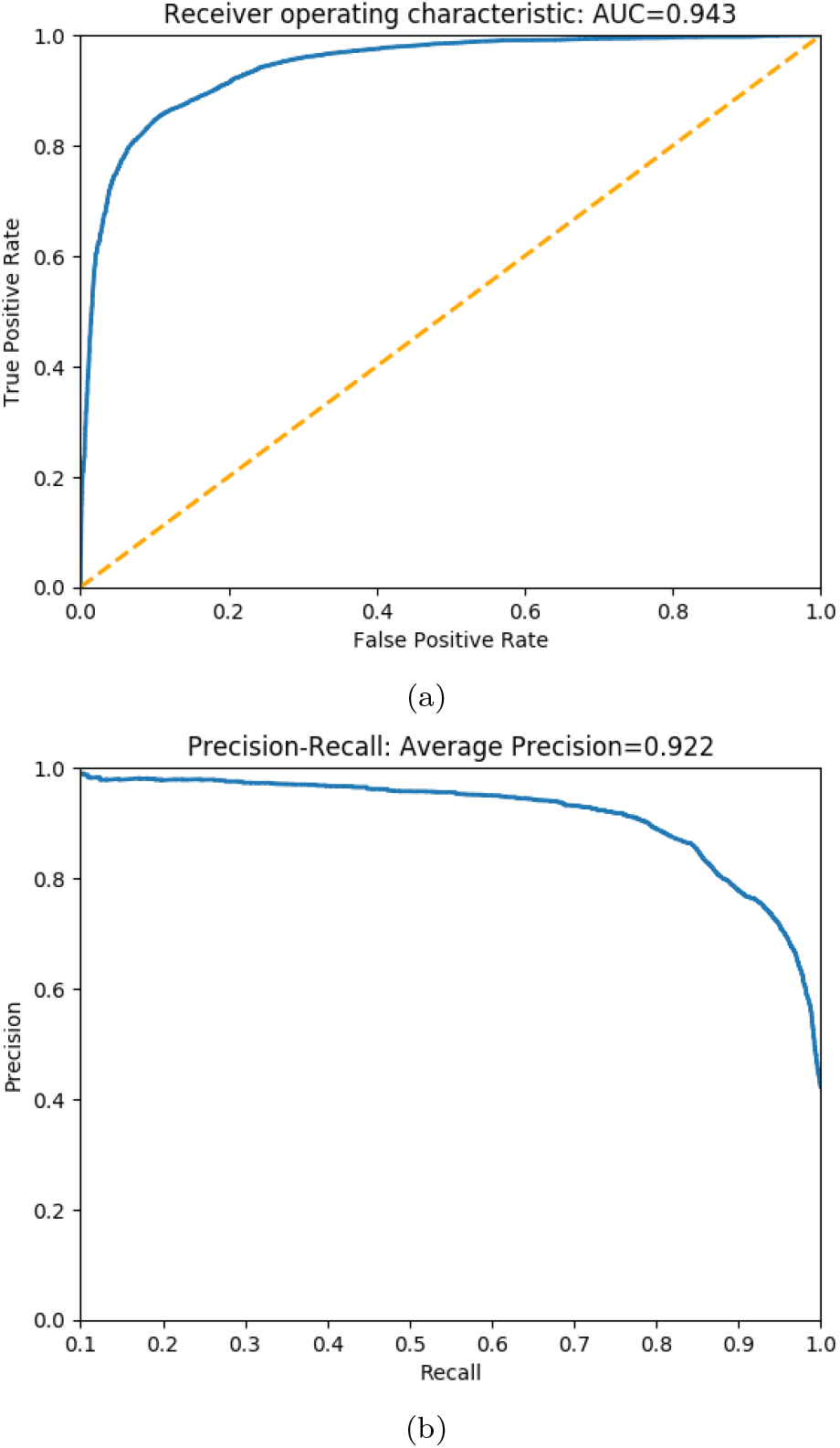
Performance of DeepPVP on a stratified-by-disease test set (25% split)

As our third evaluation, we evaluate the performance of DeepPVP using causative variants added to the ClinVar database on or after Feb 7th, 2017, while limiting our training data to all variants that have been added to ClinVar before this date. Between Feb 7th, 2017 and Feb 6th, 2018, there were 5,686 causative variants added to ClinVar, covering 1,370 diseases. 297 of these diseases were not present in our training data. This kind of evaluation allows us to estimate under more realistic conditions how well DeepPVP is able to prioritize novel variants. To evaluate DeepPVP’s performance under even more realistic conditions, we generate synthetic patient exomes by inserting a causative variant from the validation set in a randomly selected exome from the 1000 Genomes Project [24]. We remove all variants with minor allele frequency greater than 1% (using the frequencies provided by the 1000 Genomes across all populations). For each exome, we insert a causative variant from our validation set so that each exome has exactly one pathogenic variant inserted. We then assign the phenotypes associated with the causative variant in ClinVar, as well as the mode of inheritance of the disease, to the synthetic exome and consider this combination a synthetic patient. We then use DeepPVP to prioritize variants given the synthetic patient’s filtered VCF file, phenotypes, and mode of inheritance, and determine the rank at which the causative (inserted) variant is found. For comparison, we use the Exomiser version 7.2.1 released on Feb 6th, 2017 for benchmarking purposes, as well as PVP v1.1 which uses a random forest classifier trained on the same variant and phenotype data as DeepPVP. Furthermore, we compare the performance against CADD [3], DANN [35], and GWAVA [36]. Table 1 shows the results. We find that DeepPVP has a significantly improved performance compared to the original PVP, and that DeepPVP and PVP outperform Exomiser, CADD, DANN, and GWAVA.

**Table 1:** Comparison of top ranks of ClinVar variants as recovered from WES data filtered by MAF > 1% using different methods.

|  | Top hit | Top 10 hits | Total | ROC AUC |
| --- | --- | --- | --- | --- |
| DeepPVP | 4,096 (72.04%) | 4,768 (83.86%) | 5,686 | 0.95 |
| PVP | 3,619 (63.65%) | 4,076 (71.68%) | 5,686 | 0.95 |
| Exomiser | 2,910 (51.18%) | 3,608 (63.45%) | 5,686 | 0.89 |
| CADD | 1,060 (18.64%) | 2,429 (42.72%) | 5,686 | 0.94 |
| DANN | 170 (2.99%) | 1,322 (23.25%) | 5,686 | 0.90 |
| GWAVA | 63 (1.11%) | 264 (4.64%) | 5,686 | 0.66 |

Of the 5,686 “new” variants in our ClinVar evaluation set, 5,489 are in 934 genes which are associated with phenotypes. These 5,489 variants are associated with 1,289 diseases. Only 197 variants are in 74 novel genes and are associated with 89 diseases. We test the performance of DeepPVP separately on these 197 variants. DeepPVP identifies 53 of the 197 variants (27%) at rank one, and 90 variants (46%) in the first ten ranks. In comparison, Exomiser and CADD identified 26 and 13 variants at the first, and 61 and 51 variants in the top ten ranks, respectively. This evaluation demonstrates that DeepPVP can not only identify variants in known disease-associated genes but also in novel genes. While the predictive performance of DeepPVP in this evaluation is lower than in the other types of evaluation, DeepPVP still improves over established methods such as CADD and Exomiser.

Our performance results demonstrate that DeepPVP can identify causative variants with significantly higher recall at rank one and rank ten than several other methods, including the PVP system from which DeepPVP is derived. In some applications of variant prioritization, it is also important to identify causative variants quickly and with low computational costs. We therefore benchmarked the time it takes DeepPVP to process large VCF files. We used a machine equipped with 128 GB Memory and an Intel Xeon ES-2680 v3 CPU with 2.50GHz and 16 cores, using a 64-bit Ubuntu 14.04 LTS system. We selected a genome from the Personal Genome Project (PGP) [37] which contains 4,120,185 variants to benchmark DeepPVP. We prioritized variants in this genome using DeepPVP ten times and recorded the time elapsed. On average, it took DeepPVP 85 minutes to fully analyze the VCF file.

## Conclusions

DeepPVP is an easy to use and fast phenotype-based tool for prioritizing variants in personal whole exome or whole genome sequence data. DeepPVP takes a VCF file of an individual as input, together with an ontology-based description of the phenotypes observed in an individual. It then aims to identify the variants of the individual that are causative of the phenotypes observed.

Through the use of a deep neural network, an updated training and evaluation strategy, DeepPVP improves over its predecessor PVP, and further outperforms several established methods for variant prioritization, including the phenotype-based tool Exomiser [17, 18] and pathogenicity scoring algorithms such as CADD [3]. Importantly, DeepPVP shows a better performance than other methods in finding variants in novel genes, i.e., genes not previously associated with a disease phenotype, and may therefore be particularly suited for investigating variants in orphan diseases as well as variants of unknown significance in genes not yet associated with phenotypes.

We update DeepPVP in regular intervals when new training data (i.e., variants associated with diseases and phenotypes, as well as gene-phenotype associatons) becomes available. DeepPVP is freely available at https://github.com/bio-ontology-research-group/phenomenet-vp.

## Availability and requirements

- Project name: DeepPVP
- Project home page: https://github.com/bio-ontology-research-group/phenomenet-vp
- Operating system: Java virtual machine
- Programming language: Java, Groovy, Python
- Other requirements: none
- License: 4-clause BSD-style license

## List of abbreviations

### Declarations

Ethics approval and consent to participate Not applicable.

Consent for publication Not applicable.

## Availability of data and materials

All data and materials, including software developed, is freely available from https://github.com/bio-ontology-research-group/phenomenet-vp.

## Competing interests

The authors declare that they are currently applying for a patent relating to the content of the manuscript.

## Funding

This work was supported by funding from King Abdullah University of Science and Technology (KAUST) Office of Sponsored Research (OSR) under Award No. URF/1/3454-01-01 and FCC/1/1976-08-01. GVG acknowledges support from H2020-EINFRA (731075) and the National Science Foundation (IOS:1340112) as well as support from the NIHR Birmingham ECMC, NIHR Birmingham SRMRC and the NIHR Birmingham Biomedical Research Centre and the MRC HDR UK. The views expressed in this publication are those of the authors and not necessarily those of the NHS, the National Institute for Health Research, the Medical Research Council or the Department of Health.

## Author’s contributions

IB and RH conceived of the use of a deep neural network, IB and MK implemented DeepPVP, IB trained the models, performed experiments, and evaluated DeepPVP’s performance, IB, PNS, GVG, RH interpreted results and critically evaluated the performance, all authors contributed to writing the manuscript. All authors have read and approved the final version of the manuscript.

**Additional Files**

Additional file 1 — Features used to train DeepPVP

A table consisting of the features and their representation used in the training and prediction of DeepPVP.

## References

1. Eilbeck, K., Quinlan, A., Yandell, M.: Settling the score: variant prioritization and mendelian disease. Nature Reviews Genetics 18(10), 599–612 (2017). doi:10.1038/nrg.2017.52

2. Huang, Y.-F.F., Gulko, B., Siepel, A.: Fast, scalable prediction of deleterious noncoding variants from functional and population genomic data. Nature genetics 49(4), 618–624 (2017)

3. Kircher, M., Witten, D., Jain, P., Oroak, B., Cooper, G., Shendure, J.: A general framework for estimating the relative pathogenicity of human genetic variants. Nature Genetics 46(3), 310–315 (2014). doi:10.1038/ng.2892

4. Quang, D., Chen, Y., Xie, X.: Dann: a deep learning approach for annotating the pathogenicity of genetic variants. Bioinformatics 31(5), 761–763 (2015)

5. MacArthur, D.G., Tyler-Smith, C.: Loss-of-function variants in the genomes of healthy humans. Hum Mol Genet 19(R2), 125–30 (2010)

6. Robinson, P.N., Köhler, S., Oellrich, A., Project, S.M.G., Wang, K., Mungall, C.J., Lewis, S.E., Washington, N., Bauer, S., Seelow, D., Krawitz, P., Gilissen, C., Haendel, M., Smedley, D.: Improved exome prioritization of disease genes through cross-species phenotype comparison. Genome Res 24(2), 340–348 (2014). doi:10.1101/gr.160325.113

7. Boudellioua, I., Mahamad Razali, R.B., Kulmanov, M., Hashish, Y., Bajic, V.B., Goncalves-Serra, E., Schoenmakers, N., Gkoutos, G.V., Schofield, P.N., Hoehndorf, R.: Semantic prioritization of novel causative genomic variants. PLOS Computational Biology 13(4), 1–21 (2017). doi:10.1371/journal.pcbi.1005500

8. Sifrim, A., Popovic, D., Tranchevent, L.-C., Ardeshirdavani, A., Sakai, R., and Joris R Vermeesch, P.K., Aerts, J., Moor, B.D., Moreau, Y.: eXtasy: variant prioritization by genomic data fusion. Nature Methods 10, 1083–1084 (2013)

9. Singleton, M.V., Guthery, S.L., Voelkerding, K.V., Chen, K., Kennedy, B., Margraf, R.L., Durtschi, J., Eilbeck, K., Reese, M.G., Jorde, L.B., Huff, C.D., Yandell, M.: Phevor combines multiple biomedical ontologies for accurate identification of disease-causing alleles in single individuals and small nuclear families. The American Journal of Human Genetics 94(4), 599–610 (2014). doi:10.1016/j.ajhg.2014.03.010

10. de Bono, B., Hoehndorf, R., Wimalaratne, S., Gkoutos, G.V., Grenon, P.: The ricordo approach to semantic interoperability for biomedical data and models: strategy, standards and solutions. BMC Research Notes 4(1), 313 (2011)

11. Gkoutos, G.V., Green, E.C., Mallon, A.-M.M., Hancock, J.M., Davidson, D.: Using ontologies to describe mouse phenotypes. Genome biology 6(1), 5 (2005). doi:10.1186/gb-2004-6-1-r8

12. de Angelis, M.H., Nicholson, G., Selloum, M., White, J.K., Morgan, H., Ramirez-Solis, R., Sorg, T., Wells, S., Fuchs, H., Fray, M., Adams, D.J., Adams, N.C., Adler, T., Aguilar-Pimentel, A., Ali-Hadji, D., Amann, G., André, P., Atkins, S., Auburtin, A., Ayadi, A., Becker, J., Becker, L., Bedu, E., Bekeredjian, R., Birling, M.-C., Blake, A., Bottomley, J., Bowl, M.R., Brault, V., Busch, D.H., Bussell, J.N., Calzada-Wack, J., Cater, H., Champy, M.-F., Charles, P., Chevalier, C., Chiani, F., Codner, G.F., Combe, R., Cox, R., Dalloneau, E., Dierich, A., Fenza, A.D., Doe, B., Duchon, A., Eickelberg, O., Esapa, C.T., Fertak, L.E., Feigel, T., Emelyanova, I., Estabel, J., Favor, J., Flenniken, A., Gambadoro, A., Garrett, L., Gates, H., Gerdin, A.-K., Gkoutos, G., Greenaway, S., Glasl, L., Goetz, P., Cruz, I.G.D., Götz, A., Graw, J., Guimond, A., Hans, W., Hicks, G., Hölter, S.M., Höfler, H., Hancock, J.M., Hoehndorf, R., Hough, T., Houghton, R., Hurt, A., Ivandic, B., Jacobs, H., Jacquot, S., Jones, N., Karp, N.A., Katus, H.A., Kitchen, S., Klein-Rodewald, T., Klingenspor, M., Klopstock, T., Lalanne, V., Leblanc, S., Lengger, C., le Marchand, E., Ludwig, T., Lux, A., McKerlie, C., Maier, H., Mandel, J.-L., Marschall, S., Mark, M., Melvin, D.G., Meziane, H., Micklich, K., Mittelhauser, C., Monassier, L., Moulaert, D., Muller, S., Naton, B., Neff, F., Nolan, P.M., Nutter, L.M.J., Ollert, M., Pavlovic, G., Pellegata, N.S., Peter, E., Petit-Demoulière, B., Pickard, A., Podrini, C., Potter, P., Pouilly, L., Puk, O., Richardson, D., Rousseau, S., Quintanilla-Fend, L., Quwailid, M.M., Racz, I., Rathkolb, B., Riet, F., Rossant, J., Roux, M., Rozman, J., Ryder, E., Salisbury, J., Santos, L., Schäble, K.-H., Schiller, E., Schrewe, A., Schulz, H., Steinkamp, R., Simon, M., Stewart, M., Stöger, C., Stöger, T., Sun, M., Sunter, D., Teboul, L., Tilly, I., Tocchini-Valentini, G.P., Tost, M., Treise, I., Vasseur, L., Velot, E., Vogt-Weisenhorn, D., Wagner, C., Walling, A., Wattenhofer-Donze, M., Weber, B., Wendling, O., Westerberg, H., Willershäuser, M., Wolf, E., Wolter, A., Wood, J., Wurst, W., Onder Yildirim, A., Zeh, R., Zimmer, A., Zimprich, A., Holmes, C., Steel, K.P., Herault, Y., Gailus-Durner, V., Mallon, A.-M., Brown, S.D.M.: Analysis of mammalian gene function through broad-based phenotypic screens across a consortium of mouse clinics. Nature Genetics (2015)

13. Amberger, J., Bocchini, C., Hamosh, A.: A new face and new challenges for Online Mendelian Inheritance in Man (OMIM). Hum Mutat 32, 564–567 (2011)

14. Landrum, M.J., Lee, J.M., Riley, G.R., Jang, W., Rubinstein, W.S., Church, D.M., Maglott, D.R.: Clinvar: public archive of relationships among sequence variation and human phenotype. Nucleic Acids Research (2013). doi:10.1093/nar/gkt1113. http://nar.oxfordjournals.org/content/early/2013/11/14/nar.gkt1113.full.pdf+html

15. Köhler, S., Doelken, S.C., Ruef, B.J., Bauer, S., Washington, N., Westerfield, M., Gkoutos, G., Schofield, P., Smedley, D., Lewis, S.E., Robinson, P.N., Mungall, C.J.: Construction and accessibility of a cross-species phenotype ontology along with gene annotations for biomedical research. F1000Research 2 (2013). doi:10.12688/f1000research.2-30.v1

16. Rodríguez-García, M.Á., Gkoutos, G.V., Schofield, P.N., Hoehndorf, R.: Integrating phenotype ontologies with phenomenet. Journal of Biomedical Semantics 8(1), 58 (2017). doi:10.1186/s13326-017-0167-4

17. Smedley, D., Robinson, P.N.: Phenotype-driven strategies for exome prioritization of human mendelian disease genes. Genome Medicine 7(1), 1–11 (2015). doi:10.1186/s13073-015-0199-2

18. Smedley, D., Schubach, M., Jacobsen, J.O.B., Köhler, S., Zemojtel, T., Spielmann, M., Jäger, M., Hochheiser, H., Washington, N.L., McMurry, J.A., Haendel, M.A., Mungall, C.J., Lewis, S.E., Groza, T., Valentini, G., Robinson, P.N.: A Whole-Genome analysis framework for effective identification of pathogenic regulatory variants in mendelian disease. The American Journal of Human Genetics 99(3), 595–606 (2016). doi:10.1016/j.ajhg.2016.07.005

19. Lecun, Y., Bengio, Y., Hinton, G.: Deep learning. Nature 521(7553), 436–444 (2015). doi:10.1038/nature14539

20. Grimm, D.G., Azencott, C., Aicheler, F., Gieraths, U., MacArthur, D.G., Samocha, K.E., Cooper, D.N., Stenson, P.D., Daly, M.J., Smoller, J.W., Duncan, L.E., Borgwardt, K.M.: The evaluation of tools used to predict the impact of missense variants is hindered by two types of circularity. Human Mutation 36(5), 513–523 (2015). doi:10.1002/humu.22768. https://onlinelibrary.wiley.com/doi/pdf/10.1002/humu.22768

21. Cornish, A.J., David, A., Sternberg, M.J.E.: Phenorank: reducing study bias in gene prioritization through simulation. Bioinformatics, 028 (2018)

22. Danecek, P., Auton, A., Abecasis, G., Albers, C.A., Banks, E., DePristo, M.A., Handsaker, R.E., Lunter, G., Marth, G.T., Sherry, S.T., McVean, G., Durbin, R.: The variant call format and vcftools. Bioinformatics 27(15), 2156–2158 (2011). doi:10.1093/bioinformatics/btr330

23. Köhler, S., Doelken, S.C., Mungall, C.J., Bauer, S., Firth, H.V., Bailleul-Forestier, I., Black, G.C.M., Brown, D.L., Brudno, M., Campbell, J., FitzPatrick, D.R., Eppig, J.T., Jackson, A.P., Freson, K., Girdea, M., Helbig, I., Hurst, J.A., Jähn, J., Jackson, L.G., Kelly, A.M., Ledbetter, D.H., Mansour, S., Martin, C.L., Moss, C., Mumford, A., Ouwehand, W.H., Park, S.-M., Riggs, E.R., Scott, R.H., Sisodiya, S., Vooren, S.V., Wapner, R.J., Wilkie, A.O.M., Wright, C.F., Vulto-van Silfhout, A.T., Leeuw, N.d., de Vries, B.B.A., Washingthon, N.L., Smith, C.L., Westerfield, M., Schofield, P., Ruef, B.J., Gkoutos, G.V., Haendel, M., Smedley, D., Lewis, S.E., Robinson, P.N.: The human phenotype ontology project: linking molecular biology and disease through phenotype data. Nucleic Acids Res 42(D1), 966–974 (2014)

24. 1000 Genomes Project Consortium, Auton, A., Brooks, L.D., Durbin, R.M., Garrison, E.P., Kang, H.M.M., Korbel, J.O., Marchini, J.L., McCarthy, S., McVean, G.A., Abecasis, G.R.: A global reference for human genetic variation. Nature 526(7571), 68–74 (2015)

25. Li, H.: Tabix: fast retrieval of sequence features from generic tab-delimited files. Bioinformatics 27 5, 718–9 (2011)

26. Chollet, F.: Keras. GitHub (2015)

27. Abadi, M., Agarwal, A., Barham, P., Brevdo, E., Chen, Z., Citro, C., Corrado, G.S., Davis, A., Dean, J., Devin, M., Ghemawat, S., Goodfellow, I., Harp, A., Irving, G., Isard, M., Jia, Y., Jozefowicz, R., Kaiser, L., Kudlur, M., Levenberg, J., Mané, D., Monga, R., Moore, S., Murray, D., Olah, C., Schuster, M., Shlens, J., Steiner, B., Sutskever, I., Talwar, K., Tucker, P., Vanhoucke, V., Vasudevan, V., Viégas, F., Vinyals, O., Warden, P., Wattenberg, M., Wicke, M., Yu, Y., Zheng, X.: TensorFlow: Large-Scale Machine Learning on Heterogeneous Systems. Software available from tensorflow.org (2015). https://www.tensorflow.org/

28. Pedregosa, F., Varoquaux, G., Gramfort, A., Michel, V., Thirion, B., Grisel, O., Blondel, M., Prettenhofer, P., Weiss, R., Dubourg, V., Vanderplas, J., Passos, A., Cournapeau, D., Brucher, M., Perrot, M., Duchesnay, E.: Scikit-learn: Machine learning in Python. Journal of Machine Learning Research 12, 2825–2830 (2011)

29. Nair, V., Hinton, G.E.: Rectified linear units improve restricted boltzmann machines. In: Proceedings of the 27th International Conference on International Conference on Machine Learning. ICML’10, pp. 807–814. Omnipress, USA (2010). http://dl.acm.org/citation.cfm?id=3104322.3104425

30. Kingma, D.P., Ba, J.: Adam: A method for stochastic optimization. CoRR abs/1412.6980 (2014)

31. Srivastava, N., Hinton, G., Krizhevsky, A., Sutskever, I., Salakhutdinov, R.: Dropout: A simple way to prevent neural networks from overfitting. J. Mach. Learn. Res. 15(1), 1929–1958 (2014)

32. Robinson, P.N., Köhler, S., Bauer, S., Seelow, D., Horn, D., Mundlos, S.: The human phenotype ontology: a tool for annotating and analyzing human hereditary disease. Am J Hum Genet 83(5), 610–615 (2008). doi:10.1016/j.ajhg.2008.09.017

33. Smith, C.L., Goldsmith, C.-A.W., Eppig, J.T.: The mammalian phenotype ontology as a tool for annotating, analyzing and comparing phenotypic information. Genome Biol 6(1), 7 (2004). DOI:10.1186/gb-2004-6-1-r7

34. Fawcett, T.: An introduction to ROC analysis. Pattern Recogn Lett 27(8), 861–874 (2006). doi:10.1016/j.patrec.2005.10.010

35. Quang, D., Chen, Y., Xie, X.: Dann: a deep learning approach for annotating the pathogenicity of genetic variants. Bioinformatics 31(5), 761–763 (2015). doi:10.1093/bioinformatics/btu703. http://bioinformatics.oxfordjournals.org/content/31/5/761.full.pdf+html

36. Ritchie, G.R.S., Dunham, I., Zeggini, E., Flicek, P.: Functional annotation of noncoding sequence variants. Nature Methods 11, 294–296 (2014)

37. Ball, M.P., Bobe, J.R., Chou, M.F., Clegg, T., Estep, P.W., Lunshof, J.E., Vandewege, W., Zaranek, A., Church, G.M.: Harvard personal genome project: lessons from participatory public research. Genome Med 6(2), 10 (2014)

